# The phenotype of SMC6_G514R hinge mutant of *Physcomitrium patents*

**DOI:** 10.1101/2024.07.29.605668

**Authors:** Karel J. Angelis, Marcela Holá, Radka Vágnerová, Jitka Vaculíková, Jan J. Paleček

## Abstract

The Structural Maintenance of Chromosomes (SMC) complexes play roles in cohesion, condensation, replication, transcription, and DNA repair. Their cores are composed of SMC proteins with a unique structure consisting of an ATPase head, long arm, and hinge. The direct interaction of hinges leads to the formation of SMC heterodimer. A critical SMC6 mutation G551R interrupting the interaction of SMC5 and SMC6 hinges have been previously identified in *Schizosaccharomyces pombe* within a conserved motif. Using CRISPR/Cas9 directed oligonucleotide replacement, we have introduced this G to R point mutation in SMC6 of *Physcomitrium patens* (*P. patens*) at position 514 and also at position 517 of the same hinge domain. It turned out that both mutations are not toxic and do not affect the viability of established *Ppsmc6_G514R* and *Ppsmc6_G517R* lines.

Since *P. patens* mutants with entire or partial deletion of the *SMC6* gene are not viable, we compare hinge mutants with previously established mutant line with attenuated transcription by targeted binding of deactivated Cas9 nuclease (*Ppsmc6_dCas*). We show that mutation of G to R at position 514 fully prevents the interaction of SMC6 not only with SMC5, but also NSE5 and NSE6. Surprisingly, mutation of close residue 517 has no effect at all. The *Ppsmc6_G514R* line has aberrant morphology quite similar to *Ppsmc6_dCas*, though the absence of protonemata branching and formation of gametophores is incomplete. On the contrary, the *Ppsmc6_G517R* line is morphologically more or less similar to WT. Spontaneous and bleomycin-induced mutagenesis and maintenance of the number of rDNA copies in the *Ppsmc6_G514R* line is also similar to *Ppsmc6_dCas*, while *Ppsmc6_G517R* more or less mimics WT. The sensitivity of the *Ppsmc6_G514R* line to bleomycin is not as severe as that of *Ppsmc6_dCas*, and surprisingly, the *Ppsmc6_G517R* line is even less sensitive to bleomycin than WT. Moreover, both hinge mutations have no direct effect on the rate of DSB repair in dividing and differentiated cells.

The most unique feature of the hinge mutants is interference with gene targeting (GT). Whilst GT efficiency of *Ppsmc6_G517R* and *Ppsmc6_dCas* when compared to WT is only slightly or moderately reduced, it is completely abolished in *Ppsmc6_G514R*.

Based on these results, we conclude that sufficient amounts of SMC6 and its interactions are necessary for normal moss development and genome stability, such as DNA repair and rDNA maintenance. The reduced levels of SMC6 subunit, and therefore low levels of complete SMC5/6 complex, are insufficient for acute DSB repair, however, the acute DSB repair is not affected by impaired SMC6 interactions in *Ppsmc6_G514R*. In contrast, SMC6 inability to interact with SMC5 and other partners like NSE5 and NSE6 results in abolished GT, while low levels of SMC6 have only mild effect. These data underline importance of different aspects of SMC5/6, such as its levels or interactions.

## RESULTS AND DISCUSSION

### Generation and analysis of the *P. patens SMC6*-hinge mutants

The SMC5/6 of chromosome structure maintaining (SMC) complexes still plays enigmatic roles in response to DNA damage and its repair. Crucial for performing its functions is the formation and maintenance of a circular form. Circularization of SMC5 and SMC6 is achieved through the interaction of hinge regions of folded SMC subunits on one end and by bridging of ATPase heads by the NSE1,3 and kleisin NSE4 on the other. Core *SMC* and *NSE* subunits are essential genes in most organisms studied so far. For dissection of kleisin bridging of SMC5/6, we previously induced a mutant phenotype in the bryophyte *Physcomitrium patens* (*P. patens*) by attenuating transcription of the *SMC6* and *NSE4* genes by targeted binding of dCas9 (Holá *et al*., 2021).

In this study, we go further to dissect the effect of modulation of the SMC5/6 structure by uncoupling the SMC5-SMC6 interaction. The hinge region consists of two conserved interfaces, a ‘latch’ in SMC5 and a ‘hub’ in SMC6. Defined mutations in these interfaces interfere with ssDNA interaction and have severe effects on sensitivity to DNA-damage in fission yeast and reduced viability of human cells (Alt *et al*., 2017).

Critical for hinge interaction, G551R mutation has been identified in SMC6 of *Schizosaccharomyces pombe* within a PPIG*PIG motif conserved across all so far identified SMC6’s (Alt *et al*., 2017). In *P. patens SMC6* locus, this residue is at position 514. Using CRISPR/Cas9 directed oligonucleotide replacement, we introduced this *G514R* mutation as well as close mutation *G517R* (Fig. 1a and Fig.< 1b).

**Fig. 1.**
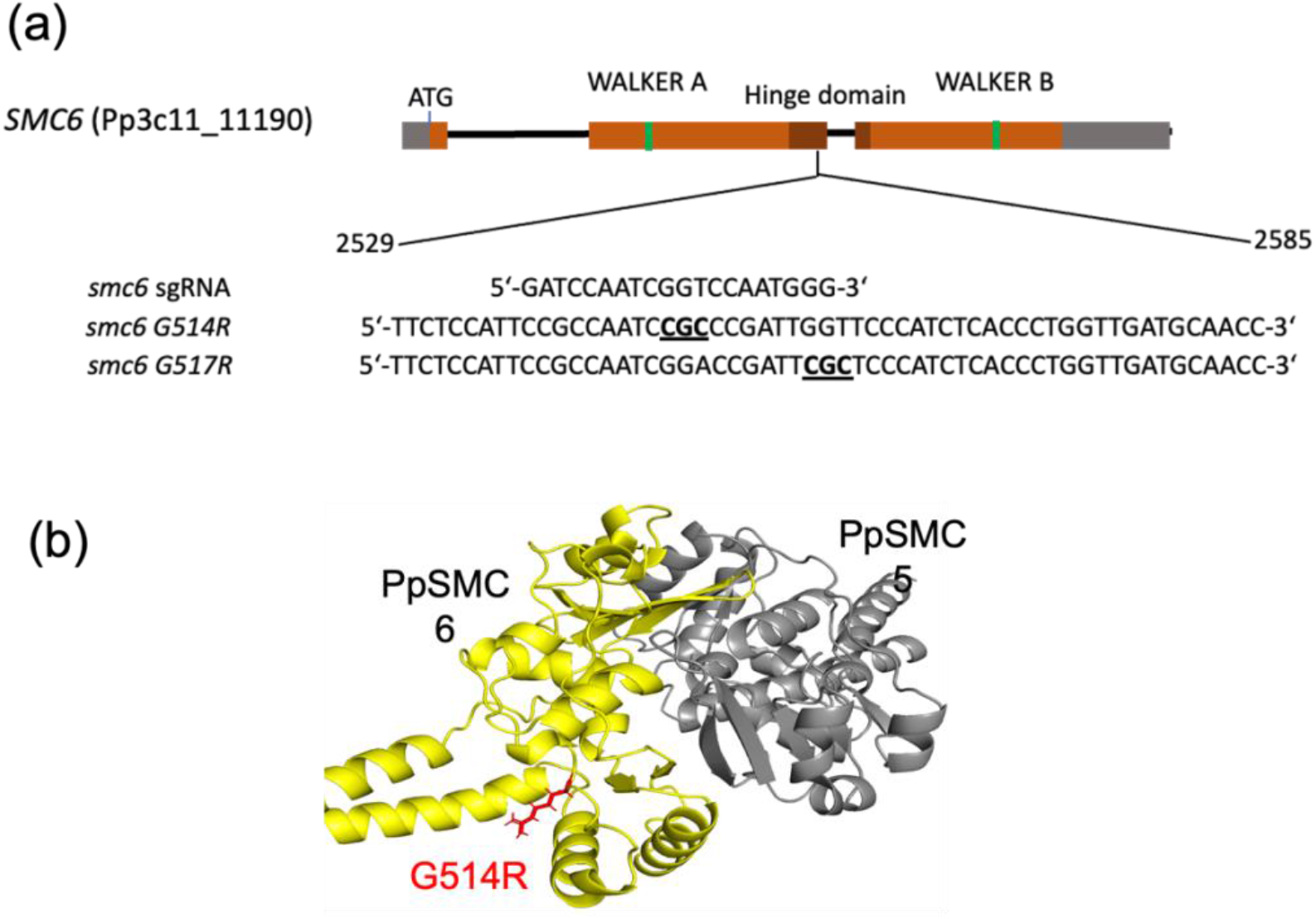
(a) Map of *P. patens SMC6* loci (Pp3c11_11190) with the site of sgRNA annealing for attenuation of *SMC6* transcription by binding ‘dead’ Cas9 nuclease (dCas9) (*Ppsmc6_dCas*), and for introduction of the donor template oligonucleotides with the point mutations G514R and G517R aided by the CRISPR/Cas9 cut. Positions of nucleotides are relative to the ATG codon, where adenine has position number 1. (b) Molecular-cartoon depiction of the *P. patens* SMC5/6 heterodimeric hinge, indicating component subdomains (yellow SMC6; grey SMC5) and localization of R, which replaced G (red) at 514.

To analyze the impact of the G514R and G517R mutations on PpSM6 interactions, we used Y2H assay (Lelkes *et al*., 2023; Vaculíková *et al*., 2024). The G514R mutation disrupted the direct interaction of SMC6 not only with SMC5, but also NSE5 and NSE6, suggesting a structural role of this residue. Surprisingly, close mutation G517R has no effect at all (Fig. 2).

**Fig. 2.**
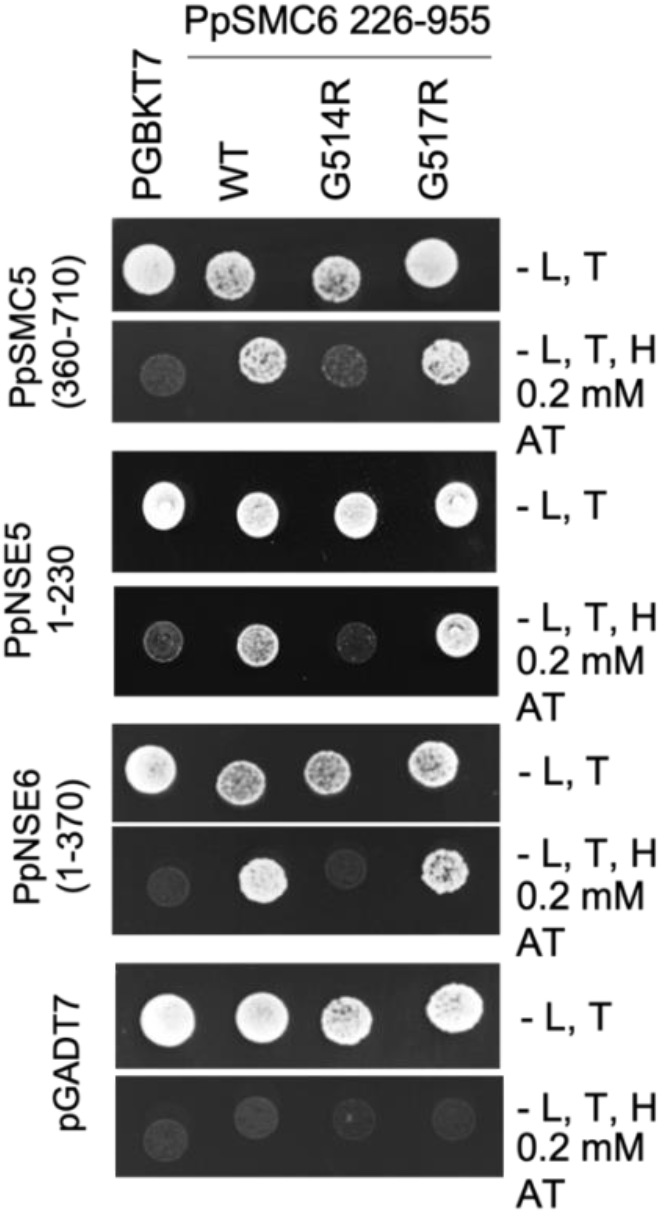
Detailed mapping of WT and mutant PpSMC6 binding to PpSMC5, PpNSE5 and PpNSE6 by Y2H. Interacting domains are indicated on the picture. While point mutation G514R interrupts the interaction of PpSMC6 with PpSMC5, PpNSE5 and PpNSE6, G517R mutation has no effect, and interactions remain the same as with WT.

### Morphology of *P. patens SMC6* hinge-mutants

Both hinge-mutations are not toxic and do not affect the viability of established *Ppsmc6_G514R* and *Ppsmc6_G517R* lines (Fig. 3a). The *Ppsmc6_G514R* line has an aberrant morphology quite similar to *Ppsmc6_dCas*, though the *Ppsmc6_G517R* line is morphologically more or less similar to WT.

**Fig. 3.**
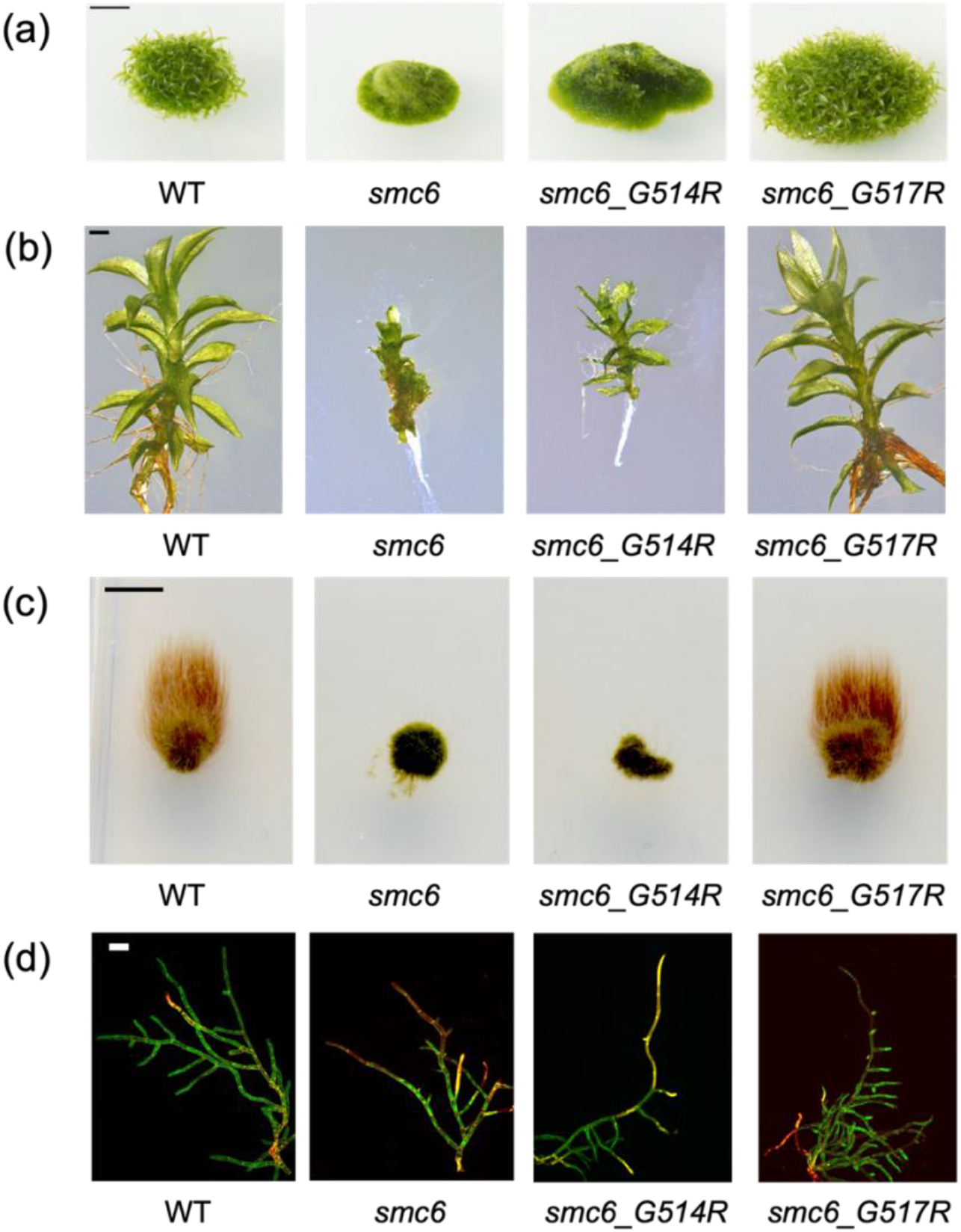
(a) Morphology of *P. patens* WT and *smc6_dCas, smc6_G514R* and *smc6_G517R* mutant lines of 1-month-old colonies grown on BCDAT medium. For simplicity, *Pp* in the names of lines on the cartoon is omitted. Scale bar 5mm. (b) Close-up view of 1-month-old gametophores. Scale bar 1mm (c) Caulonema growth after three weeks in darkness on BCDAT medium supplemented with 0.5% sucrose. Scale bar 5mm (d) Propidium iodide staining of 10-day-old protonema grown on BCDAT medium. Scale bar 100μm.

Development of gametophores in *Ppsmc6_G514R* is strongly affected, but in *Ppsmc6_dCas* nearly totally inhibited (Fig. 3a). Seldomly appearing gametophores are strongly aberrant (Fig. 3b). Nevertheless, as evident from weighting one month grown colonies shown on Fig. 3a, the viability of both lines with mutated SMC6-hinge and of WT is similar. In WT and nearly to the same extent in *Ppsmc6_G517R*, the development of juvenile gametophores is evident. In *Ppsmc6_dCas* and *Ppsmc6_G514R* mutant lines continues growth of apical cells, nevertheless they look damaged and development of gametophores is inhibited. Branching in *Ppsmc6_dCas* is nearly totally canceled by damage to the branching cells (Fig. 3d). Generally, the *Ppsmc6_G517R* line is morphologically more or less similar to WT.

### Spontaneous, but not induced mutagenesis is increased in *Ppsmc6_G514R* mutant

Spontaneous and bleomycin-induced mutagenesis and maintenance of the number of ribosomal DNA (rDNA) copies in *Ppsmc6_G514R* line is also similar to *Ppsmc6_dCas*, while *Ppsmc6_G517R* more or less mimics WT (Fig. 4 and 5).

**Fig. 4.**
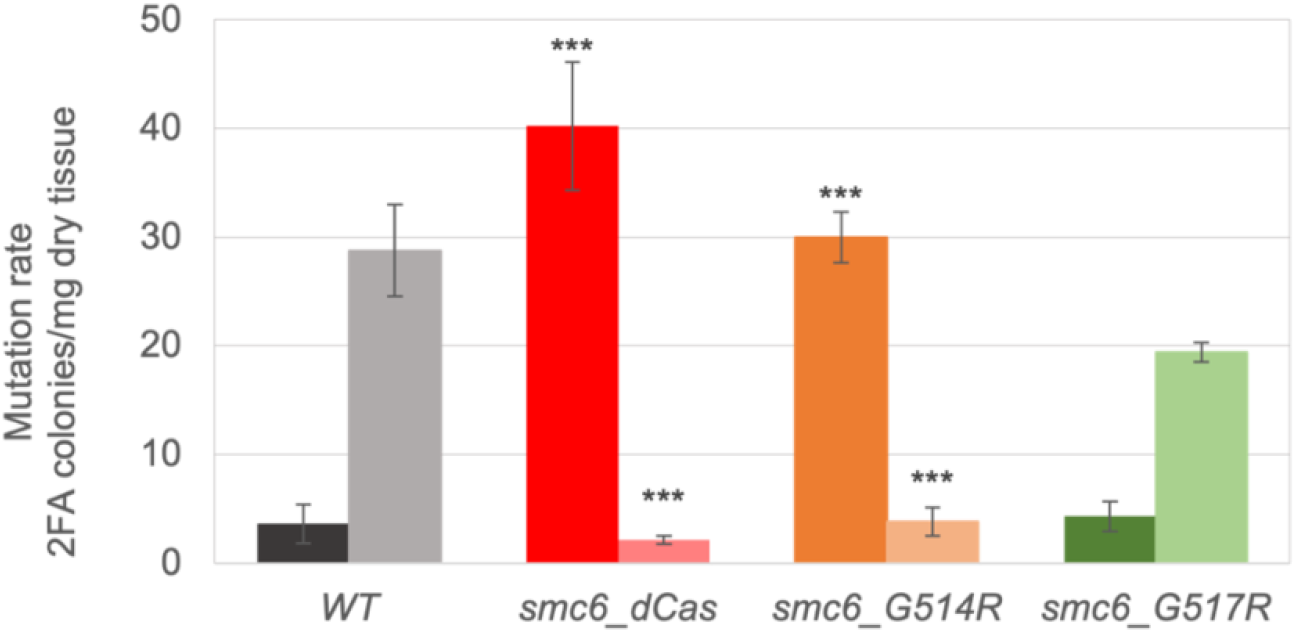
Spontaneous and bleomycin-induced mutagenesis in *APT* locus. Adenine phosphoribosyl transferase (APT) inactivated by mutation lead to the resistance to 2-fluoroadenine (2FA) (Trouiller *et al*., 2007). The 2FA surviving colonies were counted and expressed as the number of 2FA resistant colonies per mg of dry tissue. For simplicity, *Pp* in the names of lines on the graph is omitted. Student’s t-test: *P < 0.05; **P < 0.01; ***P < 0.001, and error bars represent SE.

**Fig. 5.**
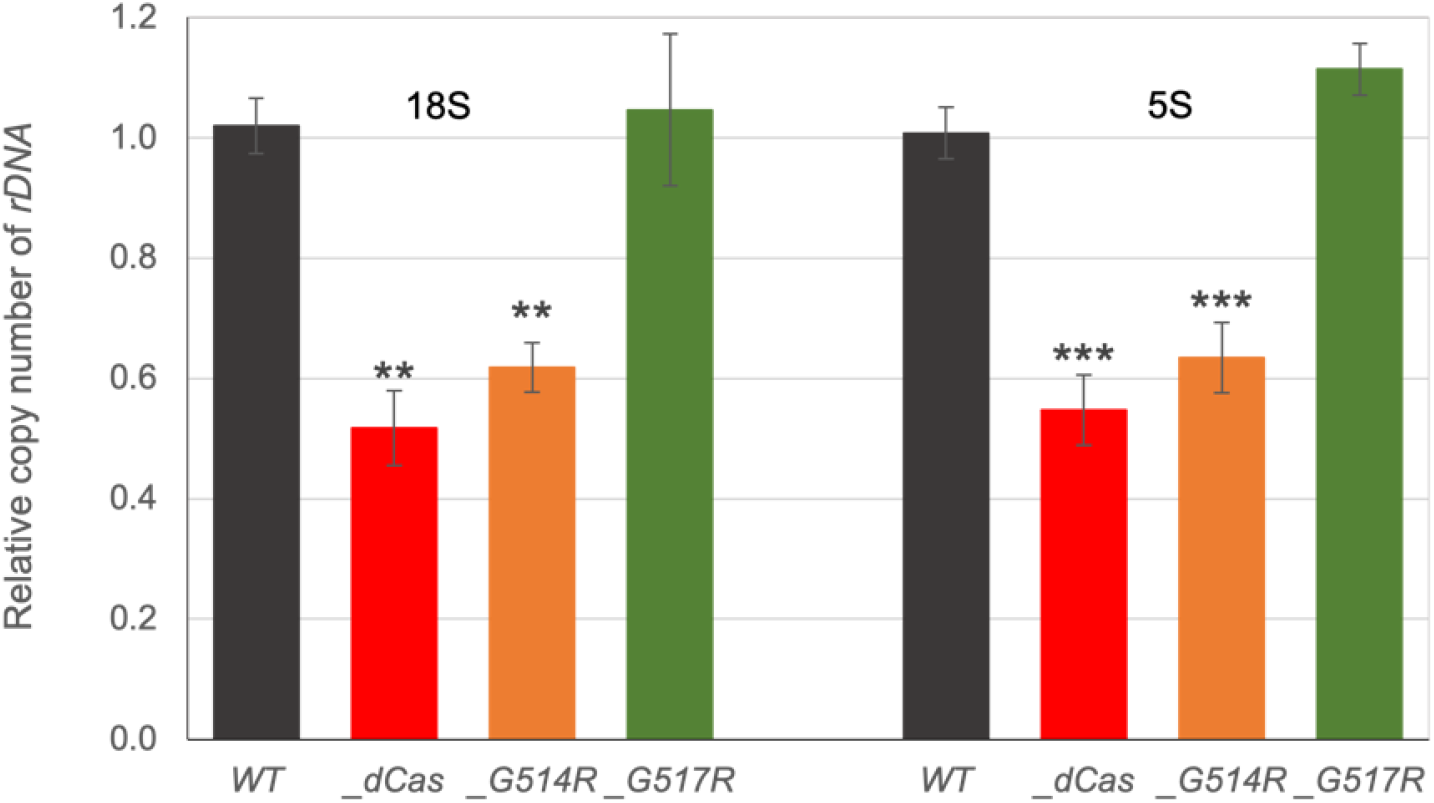
Relative copy number of 18S and 5S rDNA was measured by qPCR in *P. patens* WT (by default set to 1) and lines *Ppsmc6_dCas, Ppsmc6_G514R* and *Ppsmc6_G517R* in the graph marked for simplicity as *_dCas, _G514R* and *_G517R*. Levels of rDNA copies in *Ppsmc6_dCas* and *Ppsmc6_G514R* are significantly reduced in comparison to WT and *Ppsmc6_G517R*. Student’s t-test: *P < 0.05; **P < 0.01; ***P < 0.001, and error bars represent SE.

Spontaneous mutagenesis increases in *Ppsmc6_dCas* to 40 colonies/mg and 30 colonies/mg in *Ppsmc6_G514R*, versus only 4 colonies/mg of the dry tissue in the WT, most likely as a consequence of enhanced error-prone repair or bypass in mutated lines. In opposite to spontaneous mutagenesis, treatment with 5 μg/ml bleomycin increased the mutation rate to 28 and 19 colonies/mg of dry tissue in WT and *Ppsmc6_G517R* line, respectively, whereas in *Ppsmc6_dCas* and *Ppsmc6_G514R* lines any mutagenesis ‘induced’ by this dose of bleomycin is practically absent.

### Role of *Ppsmc6_G514R* mutant in the maintenance of genome stability

The SMC5/6 complex is actively involved in overcoming chromosome sites that are difficult to replicate. This is well documented by its role at replication fork barrier sites in the ribosomal DNA (rDNA) array of yeast (Roy *et al*., 2024). We followed the influence of hinge mutations on the rDNA maintenance by measuring a change of number of copies within arrays of 18S and 5S rDNA (Fig. 5). Levels of rDNA copies in *Ppsmc6_dCas* and *Ppsmc6_G514R* are significantly reduced, whilst line *Ppsmc6_G517R* is similar to WT, the phenotype pattern already seen in morphology and mutagenesis.

### Growth-response of *P. patens SMC6* hinge-mutants to bleomycin

Sensitivity of *Ppsmc6_G514R* line to bleomycin is not as severe as that of *Ppsmc6_dCas*. Contrary to G514R, G517R mutation has surprisingly an opposite effect and *Ppsmc6_G517R* plants are less sensitive to bleomycin and better survive bleomycin treatment than WT (Fig. 6). This suggests a flexibility of the SMC5/6 complex in orchestrating varied response to bleomycin treatment, depending on strucural modification of SMC’s and participation of subunits.

**Fig. 6.**
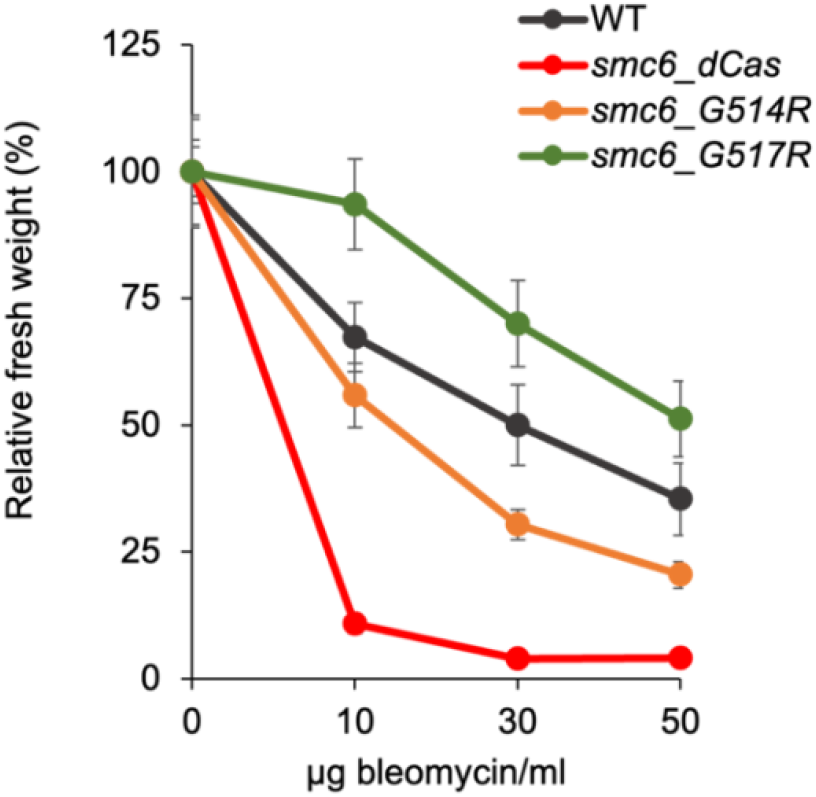
Growth-responses of *P. patens* protonemata of WT, *Ppsmc6_dCas, Ppsmc6_G514R* and *Ppsmc6_G517R* to 1 h treatment with indicated concentrations of bleomycin. For simplicity, *Pp* in the names of lines on the graph is omitted. After the treatment, the explants were incubated on a drug-free BCDAT medium under standard growth conditions for 3 weeks. Then, for each experimental point, the weight of treated plants collected from two replicas was normalized to the weight of untreated plants and plotted as relative fresh weight, which was set by default to 100. Error bars indicate SE.

### DSB repair in *P. patens SMC6* hinge-mutants is not changed

Both hinge mutations, G514R and G517R, have no effect on the rate of DSB repair in dividing (1d) cells, albeit the *Ppsmc6_dCas* line manifests a strong reduction of the rate. In differentiated (7d) cells only the line *Ppsmc6_dCas* has dramatically reduced rate of DSB repair, while *Ppsmc6_G514R* and WT rapidly repair DSB at the same pace, though much slower than in dividing cells. In addition, the *Ppsmc6_G517R* line repairs DSB even faster (Fig. 7).

**Fig. 7.**
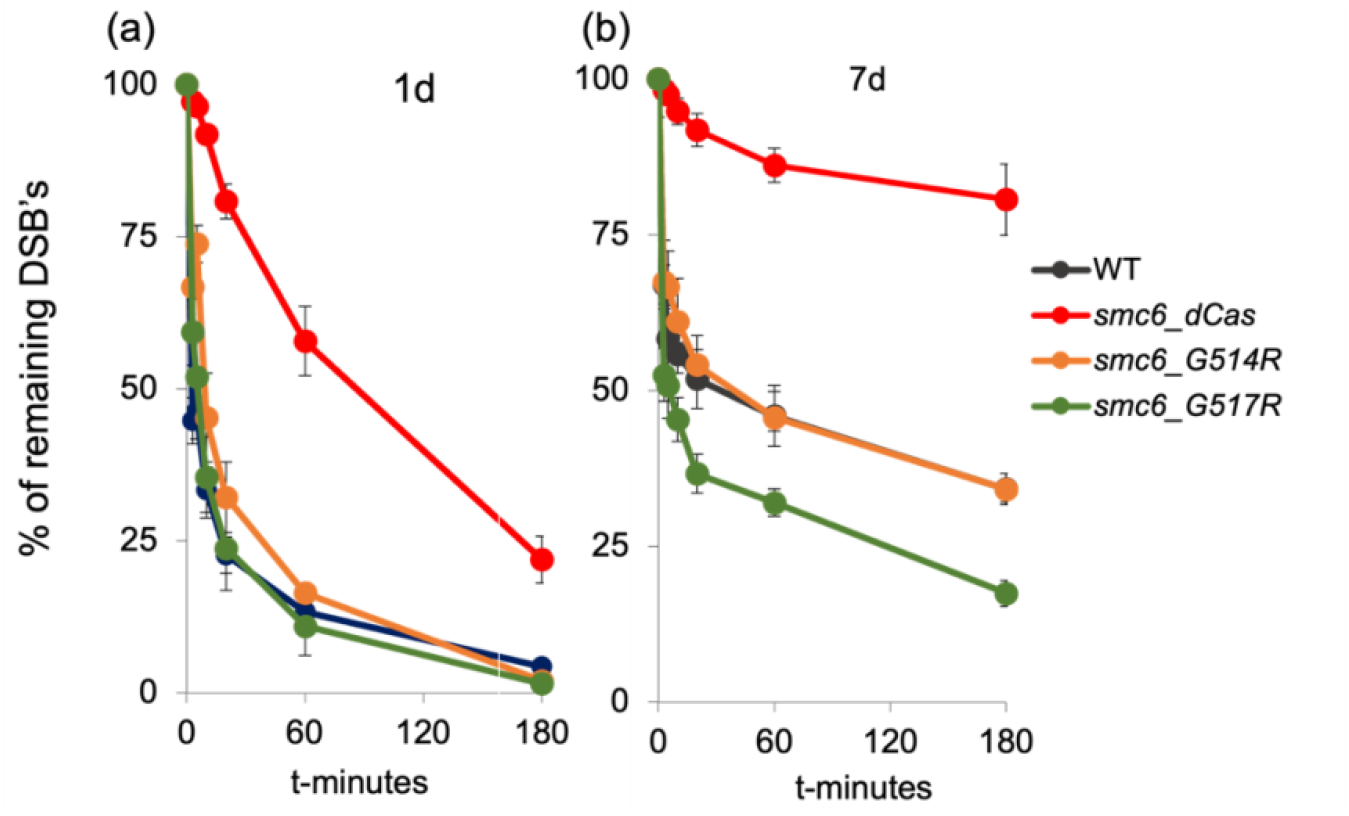
DSB repair kinetics determined by single cell gel electrophoresis (comet) assay in 1d or 7d tissue of *P. patens* WT and *Ppsmc6_dCas, Ppsmc6_G514R, Ppsmc6_G517R* mutant lines. For simplicity, *Pp* in the names of lines on the graph is omitted. Protonemata, which regenerated for 1 or 7 days after subculture, were treated with 30μg bleomycin/ml for 1 h and repair kinetics was measured as % of remaining DSBs after the 0, 3, 5, 10, 20, 60 and 180 min of repair recovery. Maximum damage is normalized as 100% at t = 0 for all lines. Error bars indicate SE.

### Gene targeting is abolished in *Ppsmc6_G514R* mutant

The most unique feature of the hinge mutants is defective gene targeting (GT) (Fig. 8).

**Fig. 8.**
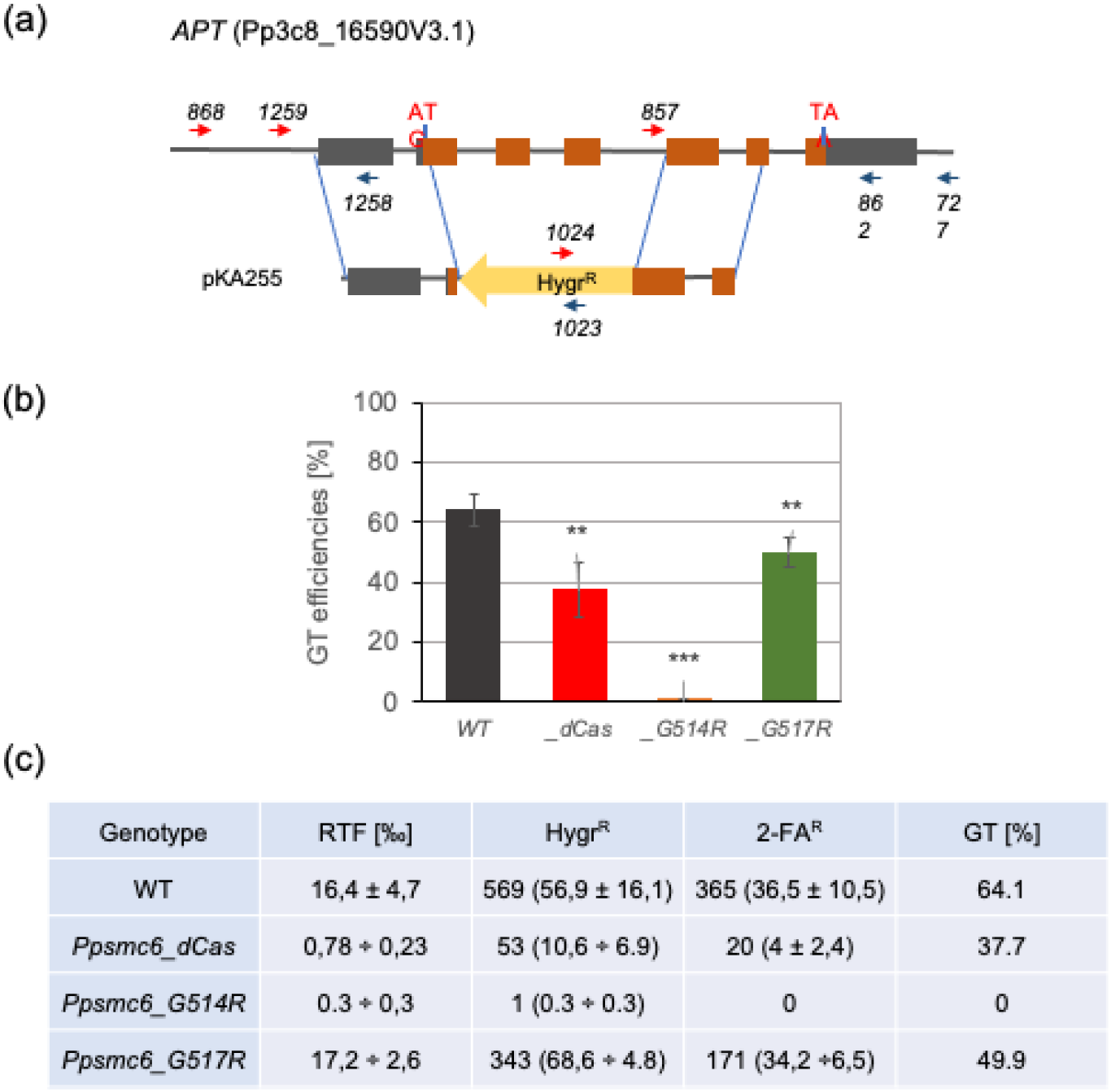
Gene targeting (GT) assay (Trouiller *et al*., 2007). (a) Schematic drawing of *P. patens APT* locus and of targeting construct pKA255. The boxes represent exons, and the gray shading indicates UTR regions. Arrows mark the position of primers used to genotype and sequence the plants by PCR (Holá *et al*., 2021). (b) GT efficiencies in *smc6_dCas, smc6_G514R* and *smc6_G517R* in the graph indicated as *_dCas, _G514R* and *_G517R*, respectively, in comparison to WT. Student’s t-test: *P < 0.05; **P < 0.01; ***P < 0.001, and error bars represent SE. (c) Comparison of transformation and GT efficiencies. Relative transformation index RTF [in ‰] is the frequency of Hygr^R^ transgenic clones in the whole regenerating protoplasts population; Hygr^R^ and 2FA^R^ indicate total and average ± SE resistant clones determined from 3 independent experiments. GT [in %] express the frequency of 2FA^R^ resistant clones among the population of Hygr^R^ transgenic clones.

Whilst GT efficiency in *Ppsmc6_G517R* is reduced by 25% and in line *Ppsmc6_dCas* by 40% when compared to WT, in *Ppsmc6_G514R* is completely abolished. Also, the transformation efficiency RTF in *Ppsmc6_G514R* is reduced nearly to zero.

## CONCLUSIONS

Based on these results, we conclude that sufficient amounts of SMC6 and its interactions are necessary for normal moss development and genome stability, such as DNA repair and rDNA maintenance. The reduced levels of SMC6 subunit, and therefore low levels of complete SMC5/6 complex, are insufficient for acute DSB repair, however, the acute DSB repair is not affected by impaired SMC6 interactions in *Ppsmc6_G514R*. In contrast, SMC6 inability to interact with SMC5 (or other partners like NSE5 and NSE6) results in abolished GT, while low levels of SMC6 have only mild effect. Loss of contact between hinges presumably interferes with the assumed binding of ssDNA and functions involving the participation of ssDNA, like during GT.

## METHODS

Methods of *P. patens* cultivation, construction and cloning of mutants and analysis of various aspects of their phenotype were in detail published in previous papers (Holá *et al*., 2021; Lelkes *et al*., 2023; Angelis *et al*., 2023; Vaculíková *et al*., 2024).

## AUTHOR CONTRIBUTIONS

Conceptualization, supervision, and funding acquisition: KJA, JJP; Data curation and formal analysis: KJA, MH, JJP, JV; Investigation: KJA, MH, RV, JJP, JV; Writing ms: KJA, JJP.

## ACKNOWLEDGEMENTS

We acknowledge skillful technical assistance of P. Rožnovská and access and use of the Imaging Facility of the Institute of Experimental Botany AS CR, supported by the MEYS CR (LM2018129 Czech-BioImaging).

## REFERENCES

Alt, A., Dang, H.Q., Wells, O.S., et al. (2017) Specialized interfaces of Smc5/6 control hinge stability and DNA association. Nat Commun, 8, 14011.

Angelis, K.J., Záveská Drábková, L., Vágnerová, R. and Holá, M. (2023) RAD51 and RAD51B Play Diverse Roles in the Repair of DNA Double Strand Breaks in Physcomitrium patens. Genes, 14, 305.

Holá, M., Vágnerová, R. and Angelis, K.J. (2021) Kleisin NSE4 of the SMC5/6 complex is necessary for DNA double strand break repair, but not for recovery from DNA damage in Physcomitrella (Physcomitrium patens). Plant Mol Biol, 107, 355–364.

Lelkes, E., Jemelková, J., Holá, M., et al. (2023) Characterization of the conserved features of the NSE6 subunit of the Physcomitrium patens SMC5 /6 complex. The Plant Journal, 115, 1084–1099.

Roy, S., Adhikary, H. and D’Amours, D. (2024) The SMC5/6 complex: folding chromosomes back into shape when genomes take a break. Nucleic Acids Research, 52, 2112–2129.

Trouiller, B., Charlot, F., Choinard, S., Schaefer, D.G. and Nogue, F. (2007) Comparison of gene targeting efficiencies in two mosses suggests that it is a conserved feature of Bryophyte transformation. Biotechnol Lett, 29, 1591–8.

Vaculíková, J., Holá, M., Králová, B., Lelkes, E., Štefanovie, B., Vágnerová, R., Angelis, K.J. and Paleček, J.J. (2024) NSE5 subunit interacts with distant regions of the SMC arms in the Physcomitrium patens SMC5/6 complex. The Plant Journal, tpj.16869.

